# A comparative study of low-pH tolerance and chitinase activity between toxigenic and non-toxigenic strains of *Vibrio cholerae*

**DOI:** 10.1101/2023.02.13.528346

**Authors:** Vijay Jayaraman, Shafqat Ali Khan, Kumar Perinbam, Isha Rakheja, Abhinav Koyamangalath Vadakkepat, Santosh Kumar Chaudhary, Asheesh Kumar Pandey, Joydeep Mitra

## Abstract

Cholera toxin, encoded by the *ctx* gene, is a key virulence factor in toxigenic *Vibrio cholerae* (*ctx*^+^) strains. However, some non-toxigenic *V. cholerae* (*ctx*^-^) strains are also pathogenic to humans and the mechanism involved in low-pH tolerance and pathogenicity in these strains remains unclear. To address this, we profiled the growth and chitinase activity in different pH of two clinical isolates of *V. cholerae*: VC20, a *ctx*^+^ strain, and WO5, a *ctx*^-^ strain. We also compared the expression level of key genes involved in pathogenesis between the strains. WO5, the non-toxigenic strain had robust growth and greater chitinase activity across a wide pH range, in comparison to VC20. Additionally, WO5 expressed higher levels of transcripts from genes implicated in host cell adhesion and virulence, namely *ompK* and *toxT*, respectively. Notably, we propose that lower *hapR* levels in WO5 contrary to VC20 is key to its low-pH tolerance. To systematically identify genes involved in low pH tolerance, we used a sequence-based homology search and found a widespread presence of low-pH adaptation modules, lysine-cadaverine, and ornithine-putrescine in multiple representative species of the *Vibrio* phylum. Furthermore, our analysis suggests that the loss of a gene encoding nitrite reductase that confers low pH tolerance is specific to *V. cholerae* and *V. mimicus*. Together, these findings reveal that the low-pH tolerance enhances the chitinase activity of the non-toxigenic strain that could help *V. cholerae* to survive the acidic environment of the stomach and readily colonize the intestine.

## Introduction

Cholera is a highly infectious diarrheal disease infecting humans, caused by the comma-shaped, gram-negative, gamma proteobacteria, *Vibrio cholerae*. The World Health Organization (WHO) estimates that *V. cholerae* causes 1.3-4.0 million cholera cases and 21,000–1,43,000 deaths worldwide (1). Multiple species in the genus *Vibrio* are pathogens with *V. cholerae* being the most notable of all (2). *V. cholerae* encompasses over 200 serogroups, however, only the serogroups O1 and O139 have been responsible for cholera epidemics and pandemics thus far (3).

*V. cholerae*, like many other *Vibrio* species is an aquatic bacteria associated with zooplankton and phytoplankton (4). The Evolution of *V. cholerae* from its free-living photosynthetic ancestor (5, 6) to a successful human pathogen has required biochemical adaptation. In the gut, the chitinase activity of the bacterium is important for colonization, where the chitinase enzyme degrades the mucin layer of the intestine and uses it as an energy source (7, 8). Chitinase secretion depends on various factors, like the type of *Vibrio* strain, pH and nutrient composition of its microenvironment (9). Further, the low pH of the stomach poses a serious threat to the survival of the bacterium. Adaptation of bacterial metabolism to low pH usually involves the prevention of formation of acidic end products of catabolism and the removal of excess protons in the cytosol (10). While various low pH adaptation mechanisms have been described in other bacteria, how *V. cholerae* strains tolerate the passage through the acidic environment of the stomach and eventually colonize the small intestine remains poorly understood.

The most studied virulence factor involved in cholera pathogenesis is the cholera toxin encoded by the lysogenic CTX phage that integrates into the bacterial genome (3). Not all pathogenic *Vibrio* strains harbour the CTX phage and hence alternative virulence mechanisms are needed for the non-toxigenic strains to establish infection. Notably, non-toxigenic strains are ancestors for the 6^th^ and 7^th^ cholera pandemic (11) besides causing local outbreaks in many parts of the world (12–14). In this study, we compared the low pH tolerance and chitinase activity of two different clinical isolates of *V. cholerae*, VC20, a toxigenic strain (15) and WO5, a non-toxigenic strain (16). WO5 is one of the strains isolated from patients during a local cholera outbreak in southern India. Infected patients showed secretory diarrhoea indistinguishable from cholera. This clinically isolated strain lacked CTX and other enterotoxins like Zot and Ace that typically causes pathogenicity in humans (16). Knowing the growth characteristics, chitinase activity and low pH tolerance are useful to understand the environment for survival and pathogenic potential in these non-toxigenic strains.

In this study, we found that WO5 in comparison to VC20 had robust growth and chitinase activity across all tested pH conditions. We observed altered gene expression profiles for key genes involved in quorum sensing, motility, and virulence in WO5 compared to VC20. Finally, using a homology-based search, we propose that the presence of lysine-cadaverine and ornithine-putrescine pathways as a key to low-pH adaptation mechanism in many species of the *Vibrio* genus. We also show that while most other *Vibrio* species have an intact gene encoding nitrite reductase, there was a clade-specific loss of this gene in *V. cholerae* and *V. mimicus* that may enhance low pH survival.

## Results and discussion

### WO5 exhibits robust growth in a broad pH range compared to VC20

To determine the effect of pH on the growth of VC20 and WO5, we profiled their growth in LB medium at the “gut-relevant” pH of 6, 7, 8, and additionally at pH 9 (**Figure 1A, B, and C**). The growth kinetics and yield of VC20 cells varied in different pH. Notably, at pH 6, we observed a prolonged lag phase (**Figure 1B**). This might correspond to the viable but non-culturable state (VBNC) that the bacteria adopt when exposed to low pH (17). However, WO5 displayed a highly similar growth profile in all pH conditions (**Figure 1C**), including pH 6 (without any lag phase). These results suggest that *V. cholerae* WO5 adapts better to changes in pH conditions compared to VC20.

**Figure 1.**
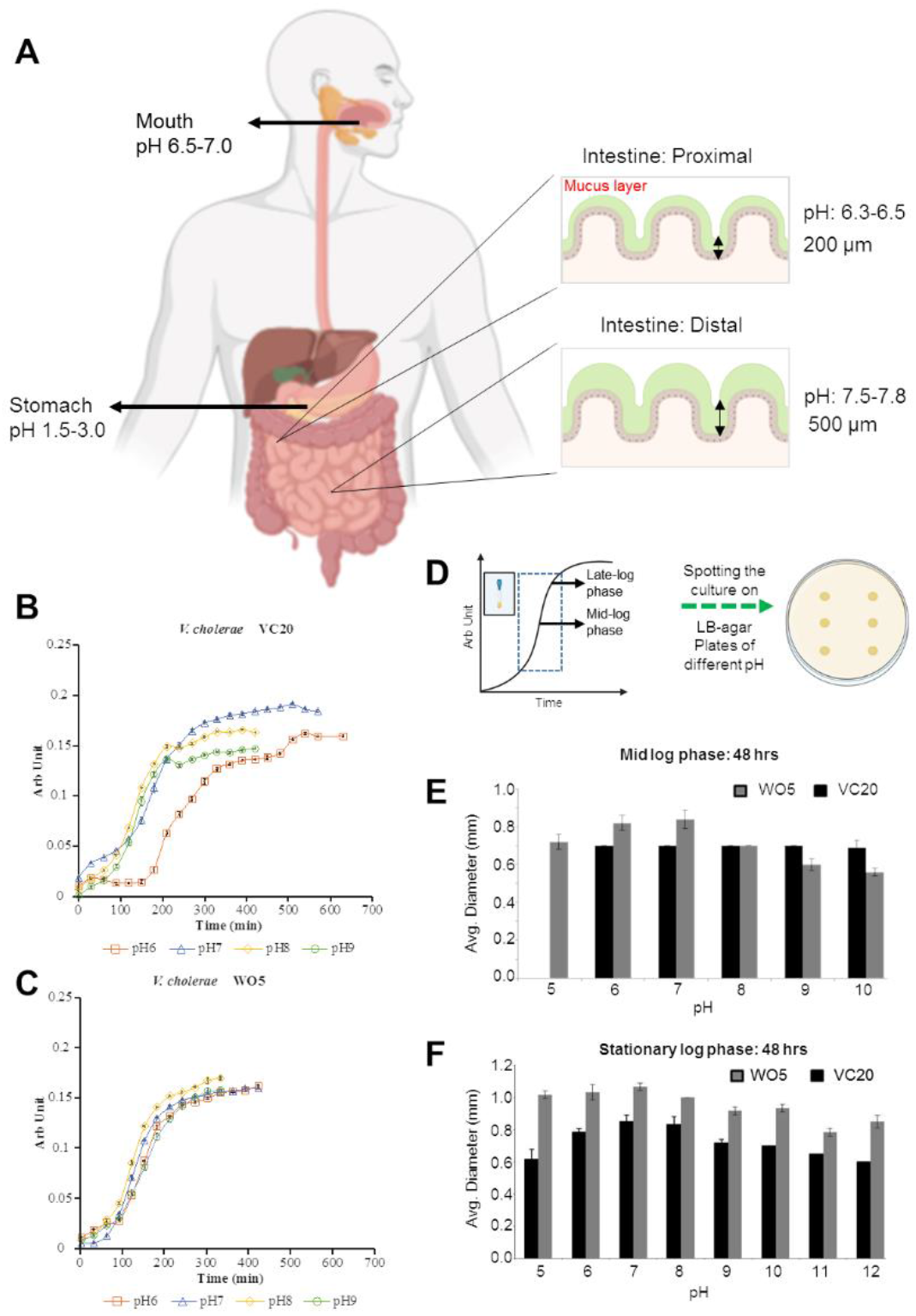
Effect of pH on the growth of *V. cholerae* strains VC20 (*ctx*^+^) and WO5 (*ctx*^-^). **A**. Cartoon showing the range of pH in different parts of the human alimentary canal and digestive system that is relevant to cholera infection. **B**. Growth curve plot of *V. cholerae* VC20 at different pH (6 to 9) in LB media. X-axis represents the main absorbance values from three independent experiments and Y-axis represents the time in minutes. Error bars indicate the standard deviations. **C**. Growth curve plot of *V. cholerae* WO5 at different pH (6 to 9) in LB media. X-axis represents the time in minutes and Y-axis represents the mean absorbance values of the culture from three independent experiments. Error bars indicate the standard deviations. **D**. Cartoon showing experimental strategy where *V. cholerae* VC20 and WO5 were grown in LB agar media and culture from mid-log and late-log phase were spotted in triplicates on LB-agar plates of different pH (5 to 10). **E**. Bar graph comparing the growth of *V. cholerae* VC20 and WO5 where mid-log phase cultures were spotted on LB-agar plates of different pH (5 to 10). X-axis represents the pH and Y-axis represents the average diameter of the spotted colonies measured at 48 hours. Error bars represent the mean ± SD of three independent experiments. **F**. Bar graph comparing the growth of *V. cholerae* VC20 and WO5, where stationary phase cultures were spotted on LB-agar plates of different pH (5 to 10). X-axis represents the pH and Y-axis represents the average diameter of the spotted colonies as measured at 48 hours. Error bars represent the mean ± SD of three independent experiments.

To understand growth-phase specific pH tolerance, cultures of VC20 and WO5 from mid-log (0.08 to 0.13 OD_600_) and late log (>0.140 OD_600_) phases were spotted in triplicate onto LB agar plates of different pH (5, 6, 7, 8, 9, 10, 11, and 12) (**Figure 1D**) and colony diameter was measured. We found that mid-log phase cultures of VC20 do not grow at pH 5, whereas WO5 does, suggesting that the strain tolerates low pH better than VC20 (**Figure 1E and S1A**). Similarly, WO5 grows better than VC20 at pH 6 and 7 (**Figure 1E**). Although mid-log cultures of VC20 and WO5 grow equally well at pH 8, VC20 grows better than WO5 at pH 9 and 10 (**Figure 1E**). Likewise, at pH 12, WO5 does not grow, whereas VC20 grows, suggesting that VC20 grows better than WO5 at alkaline pH (**Figure S1B**). When late log phase cultures were spotted, WO5 grew to a larger diameter than VC20 at all pH tested (**Figure 1F and S1C**). This result indicates that WO5 can grow to a larger mass from a stationary-phase culture across a wider pH range than VC20. From these results, it appears that WO5 can survive the low pH environment of the stomach better than VC20 and this may lead to a relatively greater bacterial load in the intestinal tract. Next, we tested if the chitinase activity is different between the strains in different pH.

### WO5 shows enhanced chitinase activity

To evaluate the effect of pH on the chitinase activity we compared the two *V. cholerae* strains, as this is key to metabolizing mucin and colonizing the intestine. For this, a semi-quantitative plate diffusion assay was carried out, by measuring the diameter of the clearing zone in the agar-colloidal chitin media plates that indicates the extent of extracellular chitinase activity (**Figure 2**). We first compared the chitinase activity of VC20 and WO5 in four different chitin-containing media formulations - LC medium (**<u>L</u>**B agar with **<u>C</u>**hitin), CC medium (LB agar with **<u>C</u>**hitin as major **<u>C</u>**arbon source), LS medium (**<u>L</u>**B agar with Chitin and **<u>S</u>**alt) and, MC medium (LB agar with **<u>M</u>**ineral and **<u>C</u>**hitin)-all containing 0.5% of chitin (**Figure 2A and B**). Both VC20 and WO5 did not show any chitinase activity in LS medium (data not shown). Both strains showed maximum chitinase activity in MC medium across all tested pH (pH 6 to 10) (**Figure 2A and 2B**), so this medium was used for all further characterizations.We hypothesize that the presence of salts and minerals in the MC medium, which more closely mirror the marine environment, enhanced the chitinase activity (18). Subsequently, we compared the time and pH-dependent kinetics of chitinase production between the strains WO5 and VC20. For this, the diffusion assay was done in MC medium at different pH (5, 6, 7, 8, 9 and 10) and the clearing zone was measured at 0, 24, 48, 72, 96 and 110 hrs. Chitinase activity of both the strains at all pH steadily increased until 72 hrs, after which there was no increase in the clearing zone diameters (data not shown for 96 and 110 hrs) (**Figure 2C and D**). Interestingly, under all the conditions tested, we observed that WO5 showed greater chitinase activity than VC20 (**Figure 2D**). For both strains, the maximum chitinase activity was at pH 6, the lowest at pH 5, and intermediate activity in the rest of the pH conditions. Next, we tested if pre-exposure of the medium to chitin had any effect on chitinase activity. We compared the activity of the strains on plates after they were grown in liquid media containing chitin **CE** or without chitin **CNE**. Surprisingly, the pre-exposure (**CE**) enhanced the chitinase activity (and probably the survival) of VC20, especially at pH 5, but it did not increase the activity of WO5 at any of the tested pHs, compared to non-exposed (**CNE**) cultures. However, WO5 had higher chitinase activity compared to VC20 throughout all the tested pH conditions, which could again aid in better intestinal colonization.

**Figure 2.**
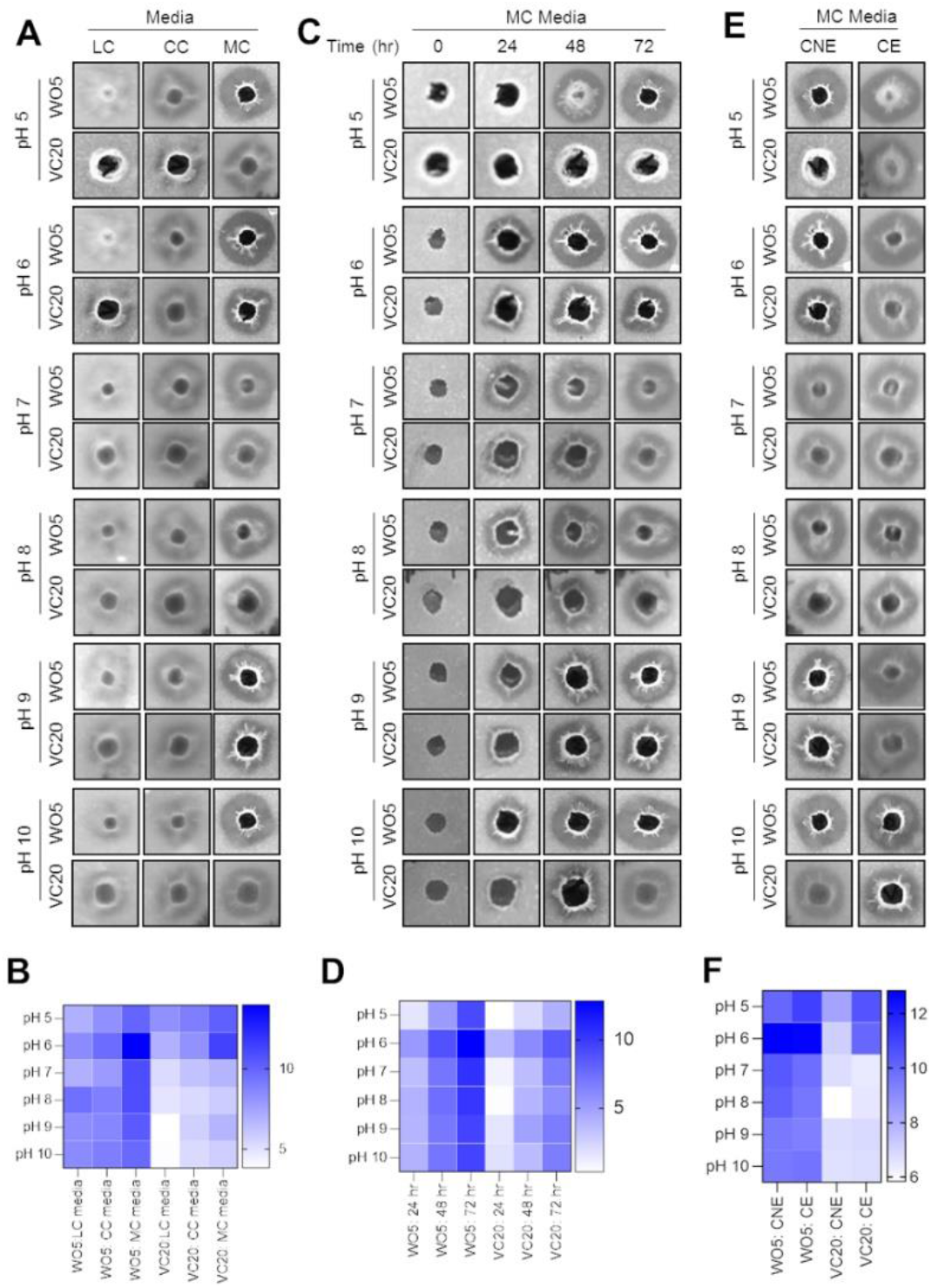
Effect of pH on the chitinase activity of *V. cholerae* VC20 (*ctx*^*+*^) and WO5 (*ctx*^*-*^). **A**. Representative images of the clearing zone at 72 hours of chitinase plate assay for *V. cholerae* VC20 and WO5 in different media conditions at a range of pH (pH 5 to 10). LC medium (**<u>L</u>**B agar with **<u>C</u>**hitin), CC medium (LB agar with **<u>C</u>**hitin as **<u>C</u>**arbon source), MC medium (LB agar with **<u>M</u>**ineral and **<u>C</u>**hitin). **B**. Heat map comparing the average diameter of the clearing zones at 72 hours of three independent chitinase plate assays for *V. cholerae* VC20 and WO5 in different media conditions at a range of pH (5 to 10). **C**. Representative images of the clearing zone of chitinase plate assay at 0, 24, 48 and 72 hours for *V. cholerae* VC20 and WO5 in MC media within a pH range of 5 to 10. **D**. Heat map comparing the average diameter of clearing zone of three independent chitinase plate assays at 0, 24, 48 and 72 hours for *V. cholerae* VC20 and WO5 in different media conditions at a range of pH (5 to 10). **E**. Representative images of clearing zone at 72 hours of chitinase plate assay for chitin non-exposed (CNE) and chitin exposed (CE) *V. cholerae* WO5 in MC medium at a range of pH (5 to 10). **F**. Heat map comparing the average diameter of clearing zone at 72 hours of three independent chitinase plate assays for chitin non-exposed and chitin exposed *V. cholerae* VC20 and WO5 in different media conditions at a range of pH (5 to 10). An arbitrary unit is used for heat map scales.

### Differential expression of HapR, ToxT and OmpK in WO5

To compare the gene expression profiles of the *V. cholerae* strains, we measured the mRNA levels (from mid-log and stationary phase) using a semi-quantitative RT-PCR of 10 judiciously selected genes (**Supplementary Table 1**) as an indicator for VC20 and WO5 adaptation during changes in pH. Crucial genes that are involved in cellular functions like cell division (*minE*) (19), quorum sensing (*hapR*), virulence gene expression (*toxT*) (20), stress response (*uspA*) (21), motility (*flgE*) (22), metabolism (*glpA*) (23), nucleoside transport and host cell adhesion (*ompK*) (24, 25), ribosomal protein (RplV), halotolerance and pH regulation (*nhaD*) (26) and hemolysin (*hlyA*) (27) were selected for this study through a comprehensive literature review. To compare the gene expression, mRNA was extracted from mid-log and stationary phase cultures of both the strains grown at different pH (**Figure 3A and Supplementary Figure 1A**). When *V. cholerae* strains were grown at pH 6 and 7, the stationary phase mRNA levels of most genes were downregulated in VC20 compared to WO5 (**Supplementary Figure 1 A)**. In the stationary growth phase, the mRNA levels of all the genes in the VC20 strain grown at pH 9 were upregulated (**Supplementary Figure 1B and 1C**) compared to WO5 highlighting that the *ctx*^*+*^ strain is better to adapt at alkaline pH, which is consistent with the growth profile (**Supplementary Figure 1B, 1E, 1F, and S1B**). The expression levels of all genes in WO5 were downregulated in pH 7 when compared to pH 6 condition (**Supplementary Figure 3E**) in the stationary phase indicating acidophilic nature of *ctx*^-^ strain, consistent with its growth profile data (**Supplementary Figure 1F**).

**Figure 3.**
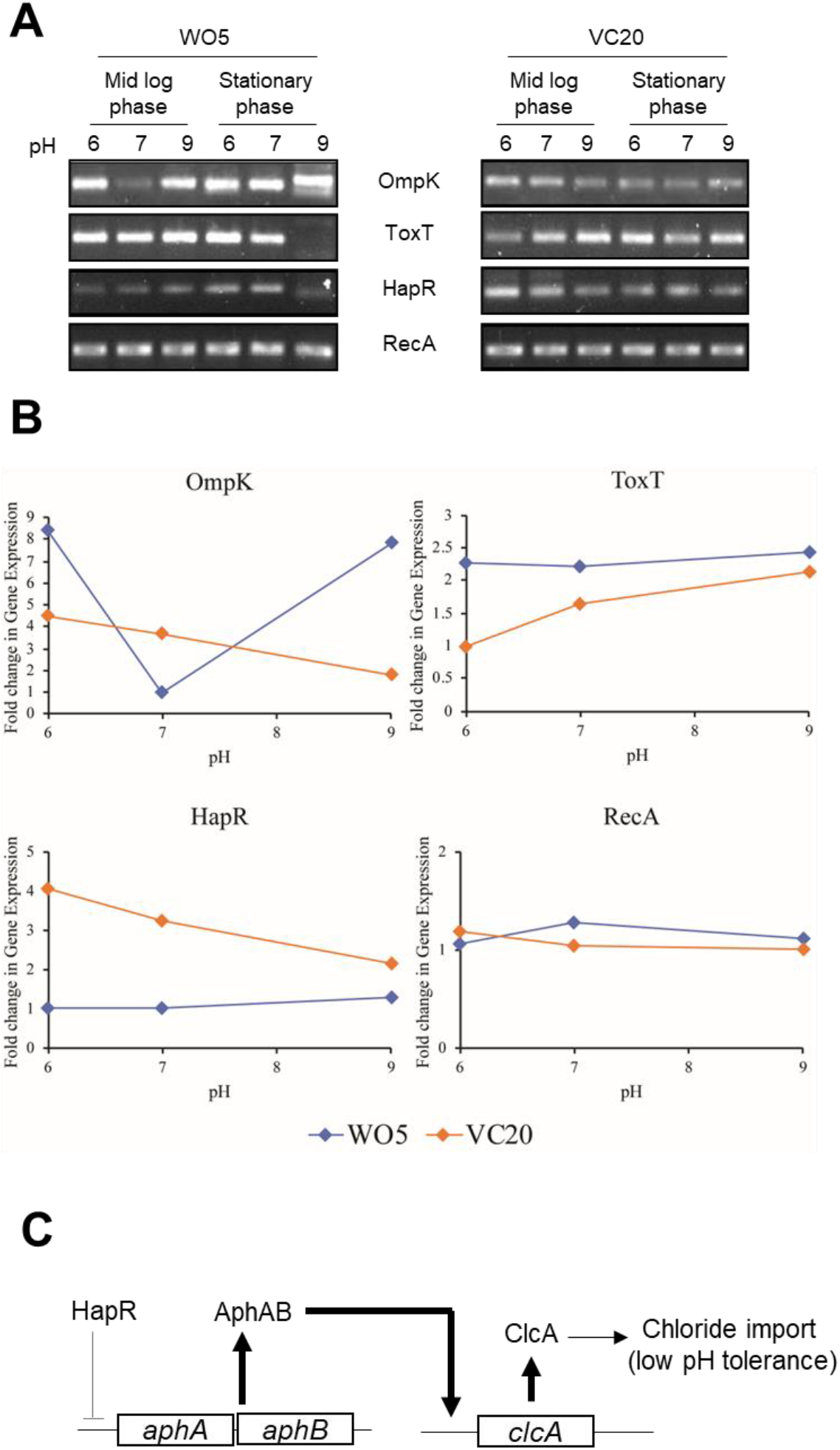
Effect of pH on the gene expression in *V. cholerae* VC20 (*ctx*^*+*^) and WO5 (*ctx*^*-*^). **A**. Agarose gel images of RT-PCR comparing expression of selected genes in *V. cholerae* VC20 and WO5 from mid-log and stationary phase at different pH (6 to 7). Only genes that had differential expression between the strains are shown here. For the complete data, see supplementary figure 2 **B**. Fold change in expression of genes between *V. cholerae* strains. **C**. HapR-based control of ClcA expression through AphAB. Lower levels of HapR in WO5 could result in enhanced expression of AphAB which in turn can directly activate the expression of *clcA* that encodes a chloride transporter. Enhanced ClcA levels in WO5 could confer low-pH tolerance in comparison to VC20.

Expression levels of many housekeeping genes like *minE, recA, flgE* were similar between the strains across all the conditions. The expression profiles of the three genes *ompK, toxT* and *hapR* were different between the two strains. In mid-log phase cultures, *toxT* and *ompK* transcripts were present at higher levels in WO5 compared to VC20 (**Figure 3A and 3B**). In *V. cholerae*, OmpK is an outer membrane protein that is implicated in the adhesion of the bacterium to its host, and nucleoside uptake (24, 25). The involvement of *ompK* in regulating chitinase activity is worth exploring in future. ToxT is a transcription factor that plays a key role in the activation of genes responsible for producing CTX and TCP (toxin-coregulated pili), which are associated with virulence. In addition, ToxT is also implicated in regulating the expression of various other gene targets (28, 29). HapR is a quorum-sensing factor involved in the repression of virulence gene expression (30). In addition to this, HapR negatively regulates the AphAB complex which is a transcriptional activator of *clcA* gene (**Figure 3C**). ClcA is a H^+^/Cl^-^ membrane protein that transports a chloride ion (in effect a negative charge) into the cytoplasm. The chloride influx is crucial to prevent inner membrane hyperpolarization caused by excess proton efflux that is typical in a low pH environment (like the stomach). Higher levels of ClcA is therefore beneficial for the growth of *V. cholerae* at low pH and detrimental at alkaline pH (31, 32). The lower expression of *hapR* could lead to higher levels of *clcA* in WO5 and this could explain both better growth at low pH ranges and lowered growth at high pH compared to VC20 (Figure 1C, E and F). Overall the results of the gene expression analysis point to the candidate genes that might explain the ability of non-toxigenic strain WO5 to tolerate low pH levels and its pathogenic mechanisms.

### Identification of genes involved in low-pH adaptation in the genus *Vibrio*

While ClcA could play an important role in preventing membrane hyperpolarization under low pH conditions, what are the mechanisms involved in the removal of excess protons that accumulate in the cytosol? Enzymes and transporters involved in the removal of excess protons are well characterized in many bacterial species. **Figure 4A** shows examples of various well characterized, low-pH adaptation modules from different bacteria. The modules generally comprise transporters and enzymes that remove protons either directly or indirectly through the production of ammonia (by deamination) or carbon dioxide (by decarboxylation) (33, 34). The presence/absence of these modules in *Vibrio* species has not been systematically investigated. To address this, we performed a homology-based search for these proteins (enzymes and transporters) in representative *Vibrio* species for which complete genome information was available. (33, 34). Guided by a robust species tree (5), we limited our search for these modules to a distinct set of *Vibrio* species (**Figure 4B**). The accession number of the query sequence used and the protein hits from the analysis are provided in **Appendix 1**.

**Figure 4.**
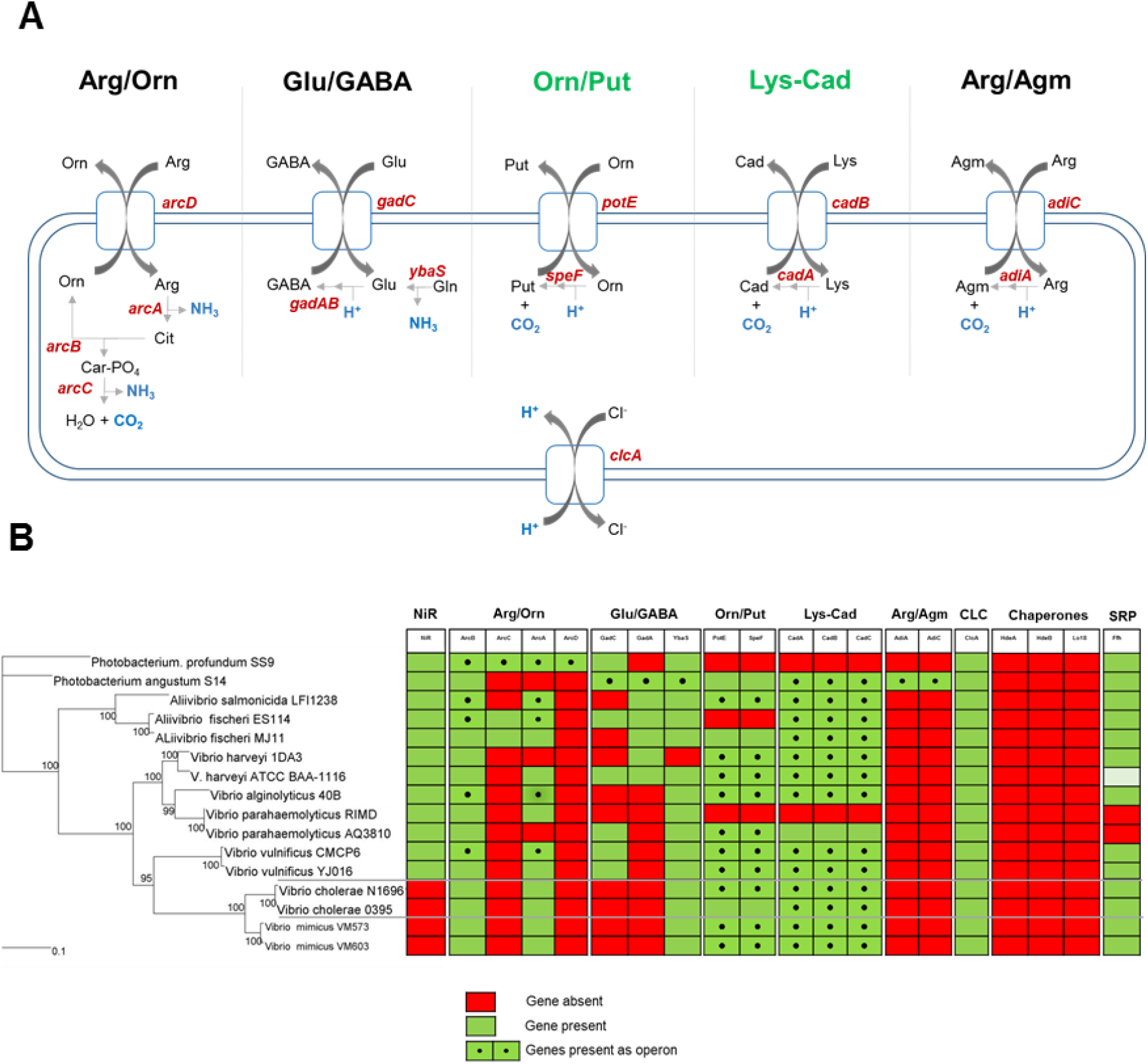
Genes involved in low-pH adaptation in the genus *Vibrio*. **A**. A schematic depiction of various low-pH adaptation modules; Arg/Orn-arginine/ornithine, Glu/GABA-glutamate/gamma-aminobutyric acid, Orn/Put-ornithine/putrescine, Lys/Cad-Lysine/cadaverine, Arg/Agm-Arginine/agmatine, *clcA*-H^+^/ Cl^-^ channel. **B**. Left-Species tree composed of various *Vibrio* strains with two *Photobacterium* strains as the outgroup. The tree is reproduced from (5). Right-Presence/absence profile of various genes in the low-pH adaptation modules listed above. Green indicates presence, red indicates absence, and green boxes with black dots in the center indicate the presence of genes in the same operon.

We found that the loss of nitrite reductase, which was recently reported to be crucial for low pH adaptation in *V. cholerae* (35), is specific to *V. cholera* and *Vibrio mimicus*. Furthermore, it was clear that lysine-cadaverine and ornithine-putrescine are the major low-pH adaptation modules which were present in all the tested *Vibrio* species (**Figure 4B**). While the lysine-cadaverine module has been previously associated with low-pH tolerance in *Vibrio* (36), the ornithine-putrescine module warrants further experimental investigation. Moreover, the genes involved in these modules were present as operons and hence will have highly correlated expression levels, which further emphasizes their direct role in low-pH adaptation. All the other modules lacked genes for one or more of the components of the pathway, without which they cannot function as low-pH counteracting modules. Importantly, the *clcA* gene that has been implicated in low-pH tolerance (31) is highly conserved in all the *Vibrio* species studied.

It was recently reported that *V. cholerae* genome despite having the gene encoding nitrate reductase, the first enzyme in nitrate metabolism (that converts nitrate to nitrite), a gene encoding the second enzyme, nitrite reductase is missing (35). This loss results in the accumulation of nitrite, which allosterically inhibits phosphofructokinase (PFK), a key enzyme in glycolysis. The inhibition of PFK lowers the glycolytic flux and prevents the accumulation of acidic end products like lactate and pyruvate, thereby counteracting a drop in cytoplasmic pH. Surprisingly, none of the *Vibrio* species used in this analysis had homologs of chaperones HdeA, HdeB, and Lo18, which are known to help fold proteins exposed to low pH (37–39). Also, except for the ATCC 1116 *Vibrio harveyi* strain, all the other strains have homologs of Ffh, which is a signal recognition particle important for membrane trafficking of these key proteins involved in low pH adaptation (40).

The low-pH tolerance repertoire of the *Vibrio* species, therefore, consists of the lysine-cadavarine and ornithine-putrescine modules that help remove excess protons from the cytosol. The ClcA transporter acts concomitantly with these proton-removal modules as an electric shunt thereby preventing inner membrane hyperpolarization. In addition to these mechanisms seen in all *Vibrio* species analyzed, the loss of nitrite reductase gene in *V. cholerae* and *V. mimicus* acts as an additional pH buffering mechanism by preventing the accumulation of acidic fermentation end products. Formation of neutral fermentation end-products like 2,3-butanediol, a characteristic feature of all *V. cholerae* El Tor biotype, can also prevent the drop in cytosolic pH (41). Variation in the regulation of expression of any of these genes in a pH-dependent manner will affect the survival and growth of both toxigenic and non-toxigenic strains in the environment and inside the human host and hence warrants detailed experimental investigation.

## Conclusion

While it is well recognized that the CTX phage-encoded cholera toxin is crucial for the pathogenicity of the toxigenic (*ctx*^+^) *V. cholerae* strains, the mechanism behind non-toxigenic (*ctx*^-^) strains that cause local cholera-like outbreaks is not clear. In this study, we compared the pH tolerance of a toxigenic and non-toxigenic cholera strain in both liquid cultures and solid growth media. We found that the non-toxigenic strain (WO5) survives and tolerates a broader pH range than the toxigenic strain (VC20). Our analyses of these strains also identified differences in the expression profiles of key pathogenic genes. The phenotypic differences that we observed between the strains could be caused by changes in the expression of some of these regulatory genes. We suggest that besides the toxin-based mechanism, greater chitinase activity and adaptation to growth in a broader pH range, relevant to the human gut, could be a key to the environmental survival of non-toxigenic *Vibrio* strains. Finally, we identified the general low-pH tolerance mechanisms in *Vibrio* species and additional mechanisms that are specific to human pathogens, *V. cholera* and *V. mimicus*. We propose that the protein sequences and gene regulatory regions of enzymes and transporters, implicated in low pH tolerance in *Vibrio* species, identified in this study can be used for systematic comparison of the two *Vibrio* strains once the whole genome sequences are available.

## Materials and Methods

### Bacterial Strains and Chemicals

For all experiments, WO5 and VC20 were used: VC20 (15) is an O1 El Tor Inaba cholera toxin *ctx*^*+*^ strain and WO5 (16) is an O1 El Tor Inaba *ctx*^*-*^ strain. Gammaproteobacteria *Escherichia coli* DH5α was used as a control. For media preparation agar, tryptone and yeast extract were purchased from HiMedia (Mumbai, India) and crab shell chitin from Sigma (cat no 417955) was used.

### Growth Kinetics or Growth Curve

For growth kinetics studies, a single colony of *V. cholerae* VC20, WO5 or *E. coli* DH5α was inoculated in 5 ml of Luria-Bertani (LB) broth at pH 5, 6, 7, 8, 9 and 10 and grown overnight at 37^°^C at 200 rpm. The pH of the growth media was adjusted using 0.1M NaOH or HCl. Growth curves for *V. cholerae* VC20, WO5 and *E. coli* DH5α were done in 250 ml conical flasks containing 100 ml LB medium of specified pH, inoculated with a 1% starter culture (v/v) of the respective strain from a saturated overnight culture. The bacterial cell density was measured in units of optical density by a Beckman model DU spectrophotometer (Beckman Instruments, Inc., Fullerton, California, U.S.A.) with 1.0 cm path length at 600 nm at every 30 minutes. To take measurements, a 1 ml sample of culture was removed aseptically from the culture flask and the blank was set in the spectrophotometer using LB medium of the same pH. Zero-hour readings were taken immediately after inoculation. All the growth curves were made in triplicates and average values were used for plotting.

### Growth assay

1.5% LB agar plates of pH 5, 5.2, 5.4, 5.6, 5.8, 6, 7, 8, 9, 10, 11 and 12 were prepared. One *μ*L of the overnight grown culture of *V. cholerae* VC20, WO5 and *E. coli* DH5α at pH 7 was spotted onto the LB agar plates of specified pH in triplicate. While spotting, the cultures were placed at ample distances to avoid the merging of colonies. The plates were allowed to dry in the laminar flow hood for 30 minutes and then incubated at 37^°^C. To take measurements, the diameter of every colony was measured every 24 hours. The experiment was done in triplicate and average values of diameter were plotted.

### Preparation of Colloidal Chitin

Colloidal chitin was prepared based on a previously published protocol (42). Briefly, 30 grams of chitin was mixed with 400 ml of concentrated ice-cold hydrochloric acid and stirred for 24 h at 4°C. The suspension was then washed with Milli-Q water until it reached a pH of 7. The chitin percentage (w/v) in the colloidal chitin suspension was determined by drying 1 ml of suspension at 37°C and then weighing it. This is done in triplicate. For the preparation of the growth medium, a final concentration of 0.5% colloidal chitin was used, as it gave the best clearing zone during standardization.

### Preparation of Colloidal Chitin LB Agar Media Plates

To measure chitinase activity of *V. cholerae*, four different LB agar-colloidal chitin media were prepared at pH 5, 6, 7, 8, 9 and 10. - LC (**<u>L</u>**B agar with **<u>C</u>**hitin) was prepared with 0.5% Yeast extract, 1.0% Tryptone, 1.0% NaCl, 1.5% Agar and 0.5% Colloidal chitin; CC media (LB agar with **<u>C</u>**hitin as major **<u>C</u>**arbon source) was prepared with 1.0% Tryptone, 1.0% NaCl, 1.5% Agar and 0.5% Colloidal chitin; LS media (**<u>L</u>**B agar with Chitin and **<u>S</u>**alt) was prepared with 0.5% Yeast extract, 1.0% Tryptone, 3.5% NaCl, 1.5% Agar and 0.5% Colloidal chitin; MC medium (LB agar with **<u>M</u>**ineral and **<u>C</u>**hitin) was prepared with 0.05% Yeast Extract, 0.2% di-Potassium hydrogen Phosphate, 0.1% Potassium dihydrogen Phosphate, 0.07% Magnesium Sulfate pentahydrate, 1.05% Sodium chloride, 0.05% Potassium chloride, 0.01% Calcium chloride, 1.5% Agar, 0.5% Colloidal chitin (42). For all four media, pH was set before autoclaving and then poured into the plates. After the media solidified, 9 holes of 5 mm diameter were aseptically punched at an equal distance using cork bores and plates were kept at 37 °C overnight to check for contamination before starting chitinase diffusion assay.

### Chitinase diffusion assay

Overnight cultures of *V. cholerae* WO5 and VC20 were first diluted to the same optical density, to ensure an equal number of cells per unit volume of culture. Five µl of diluted culture, containing an equal number of cells was inoculated in triplicates into the holes made in LB agar colloidal chitin medium plates. Thereafter, plates were kept upright for 1 hour in a laminar hood, to allow the liquid culture to get absorbed, and then were kept inverted at 37°C for 72 hours, with observations at every 24-hour time interval. Chitinase activity was assessed by measuring the diameter of the clearing zone formed due to the production of chitinase, which degrades the chitin in the agar media plates. For each clearing zone, maximum and minimum diameters were measured using a measuring scale. The experiment was performed in triplicate and average values were plotted. GraphPad Prism version 8.4.3 was used to make the heat maps.

### RNA Preparation, RT-PCR

RNA from *V. cholerae* VC20 and WO5 cells was prepared using RNeasy Mini Kit from Qiagen (Qiagen, Hilden, Germany). cDNA was prepared from an equal amount of RNA by reverse transcription using NEB mMuLV reverse transcriptase (New England Biolabs, MA, USA). The list of primers used for amplification is shown in supplementary table 2 (Sigma Aldrich, St. Louis, MO, USA). RecA was used to normalize the amount of cDNA while conducting RT PCR to measure expression levels of ten *V. cholerae* genes. PCR products were resolved on 1.5% agarose (HiMedia, Mumbai, India), formaldehyde (HiMedia, Mumbai, India) denaturing gel electrophoresis, and intensity of bands was calculated using Carestream GL21 Pro ROI analysis (Carestream Health, USA). GraphPad Prism version 8.4.3 was used to make the heat maps.

### Gene gain/loss analysis

Protein sequences of genes that have been established to be involved in low pH adaptation were collected from UniProt (43) and used as query sequences in the BLAST-P (44, 45). The homology search was done using default blast-p parameters. The search was restricted to 16 *Vibrio* species used in the construction of the species tree. An e-value of 1e-30, sequence coverage of 90 %, and sequence identity of 40 % were used as threshold values for considering a hit to be included as present in an organism.

## Acknowledgement

I.R., S.A.K., K.P., V.J., A.K.P, A.K.V and S.K.C. were supported by fellowships and M.Tech. project grants from the Department of Biotechnology, Government of India. T. Ramamurthy (National Institute of Cholera and Enteric Diseases, Kolkata, India) provided *Vibrio cholerae* strains VC20 and WO5. The authors recognize the initial contributions of former lab Mitra members, Mahendra Kumar Verma (American University School of Medicine, Aruba), Ganapathy G. (Indian Institute of Food Processing Technology Thiruvarur, India) and Sahil Ahmed (Institute of Microbial Technology, Chandigarh, India).

## Figures

**Supplementary figure 1.**
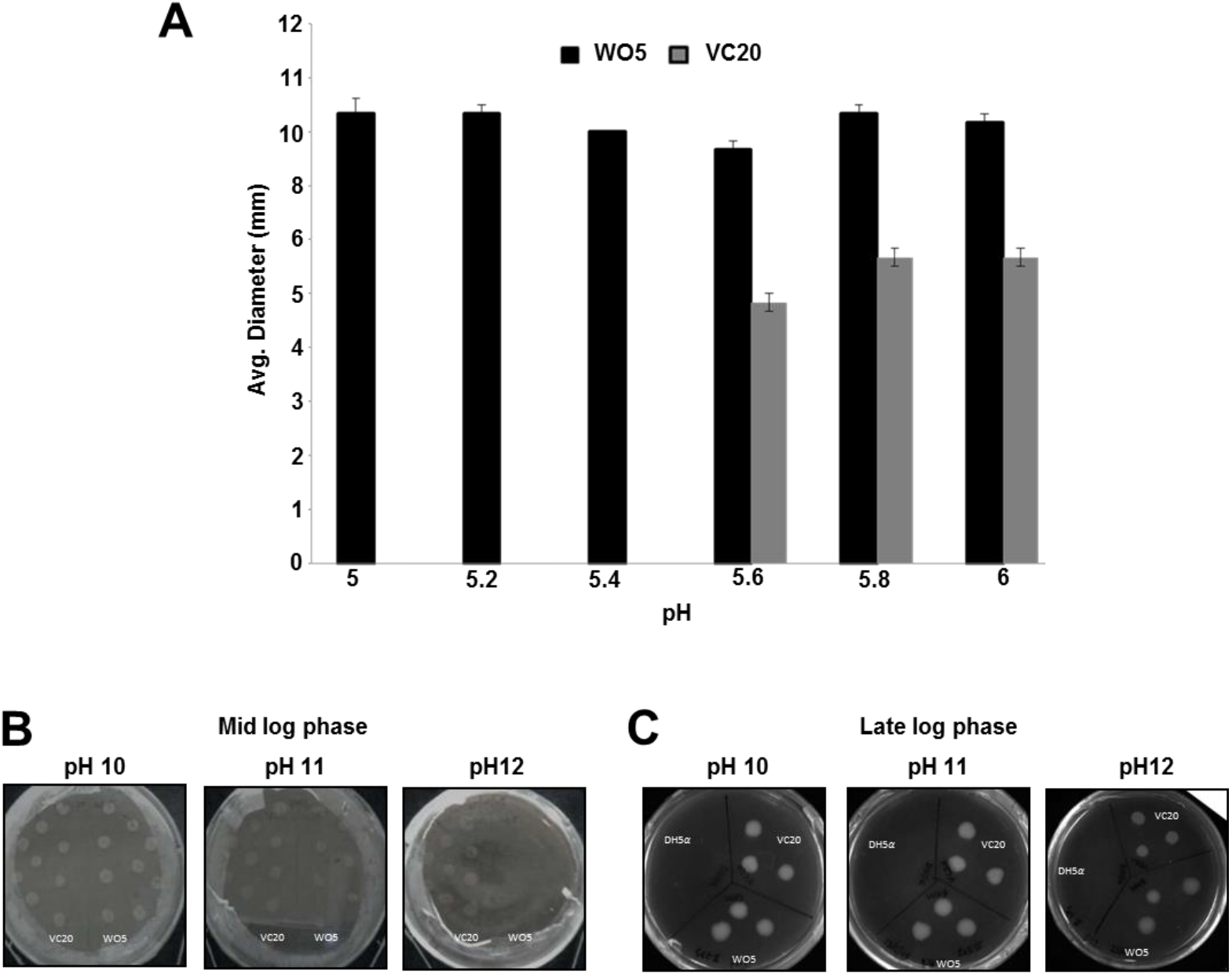
Growth profile comparison of *V. cholerae* VC20 (*ctx* +) and WO5 (*ctx* -) strains at different pH. **A**. The growth of *V. cholerae* strains VC20 and WO5 was compared at a specific pH range (5-6) by measuring the diameter (in mm) of the colonies to plot this bar graph. The average of three different experiments and standard deviations in diameter measurements were used for plotting the bar graph and error bars respectively. **B**. Mid-log phase culture of *V. cholerae* VC 20 and WO5 strains were freshly spotted side by side on a pH adjusted 2% LB agar plate to measure the growth. **C**. Stationary phase culture of *Escherichia coli* DH5*α, V. cholerae* VC 20 and WO5 strains were freshly spotted side by side on a pH adjusted 2% LB agar plate to measure the growth.

**Supplementary figure 2.**
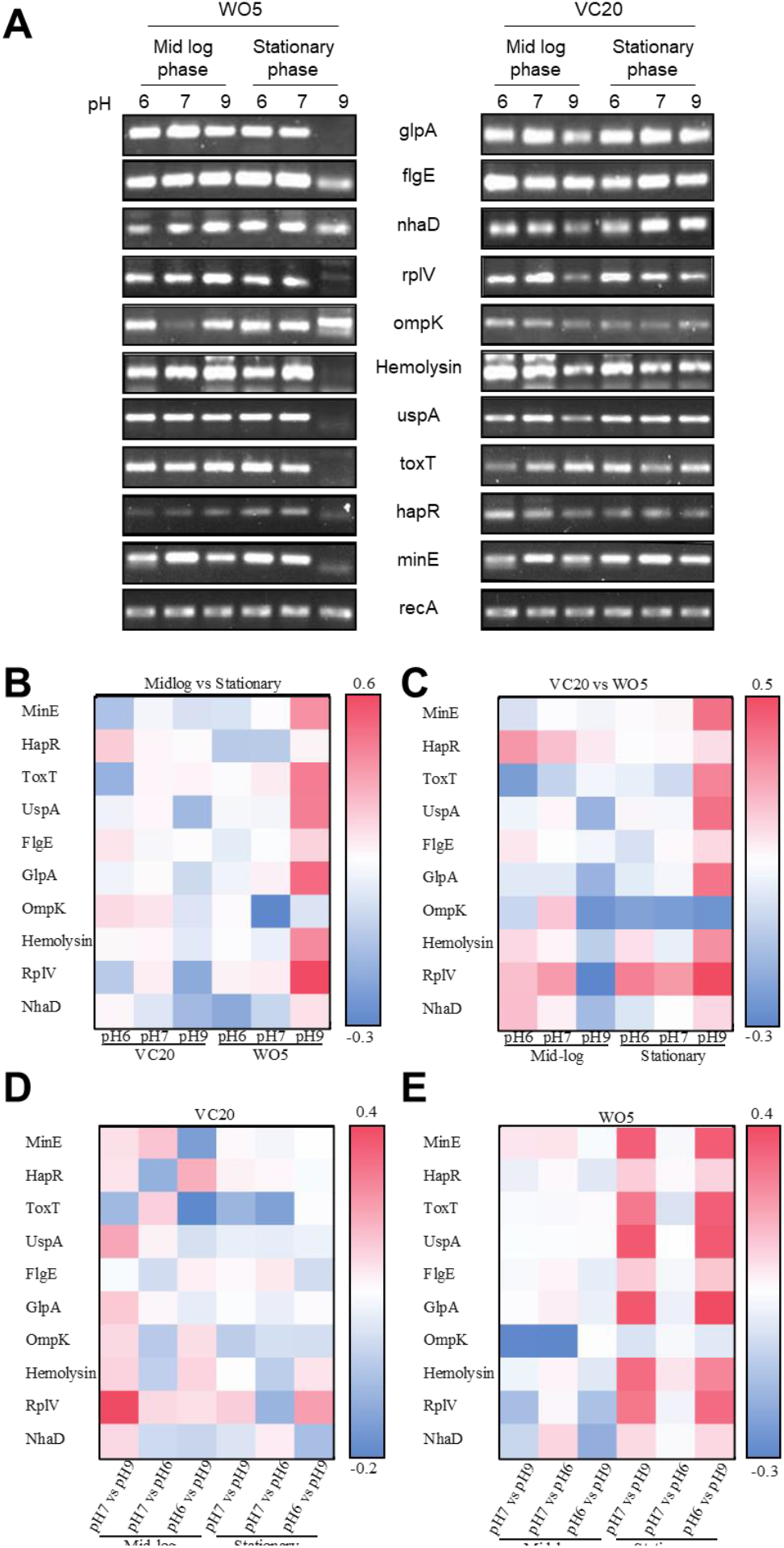
Figure 3. Effect of pH on the gene expression in *V. cholerae* VC20 (*ctx*^*+*^) and WO5 (*ctx*^*-*^). **A**. Agarose gel images of RT-PCR comparing expression of selected genes in *V. cholerae* VC20 and WO5 from mid-log and stationary phase at different pH (6 to 7). **B**. Heat map showing fold change in gene expression between mid-log and stationary phage cultures of VC20 and WO5 at different pH’s **C**. Heat map showing fold change in gene expression between VC20 and WO5 at different pH and growth phases **D**. Heat map showing fold change in gene expression in VC20 when grown in different pH **E**. Heat map showing fold change in gene expression in WO5 when grown in different pH

**Supplementary Table 1.** Genes used in this study for conducting gene expression analysis.

| S. No | Gene name | Function | NCBI gene ID |
| --- | --- | --- | --- |
| 1 | *recA* | Involved in recombination. Used as an internal control gene. | 69720701 |
| 2 | *minE* | Cell division topological specificity factor. | 69719410 |
| 3 | *hapR* | Virulence-stimulating gene, regulation of quorum sensing, low-pH adaptation | 2831115 |
| 4 | *toxT* | Essential for colonization; virulence activator gene | 2614505 |
| 5 | *uspA* | Universal stress protein | 2615765 |
| 6 | *flgE* | Flagellar hook protein | 2613237 |
| 7 | *glpA* | Glycerol-3-phosphate dehydrogenase | 2611906 |
| 8 | *ompK* | Outer membrane protein that serves as a receptor for broad-host-range Vibriophage KVP40. | 11461617 |
| 9 | Hemolysin | Causes cytolysis by forming heptameric pores in target host membranes. | 2614150 |
| 10 | *rplV* | 50S ribosomal subunit protein L22 | 2615608 |
| 11 | *nhaD* | Na<sup>+</sup>/H<sup>+</sup> antiporter; helps *Vibrios* survive in extreme salt concentrations | 2612475 |

**Supplementary Table 2.**
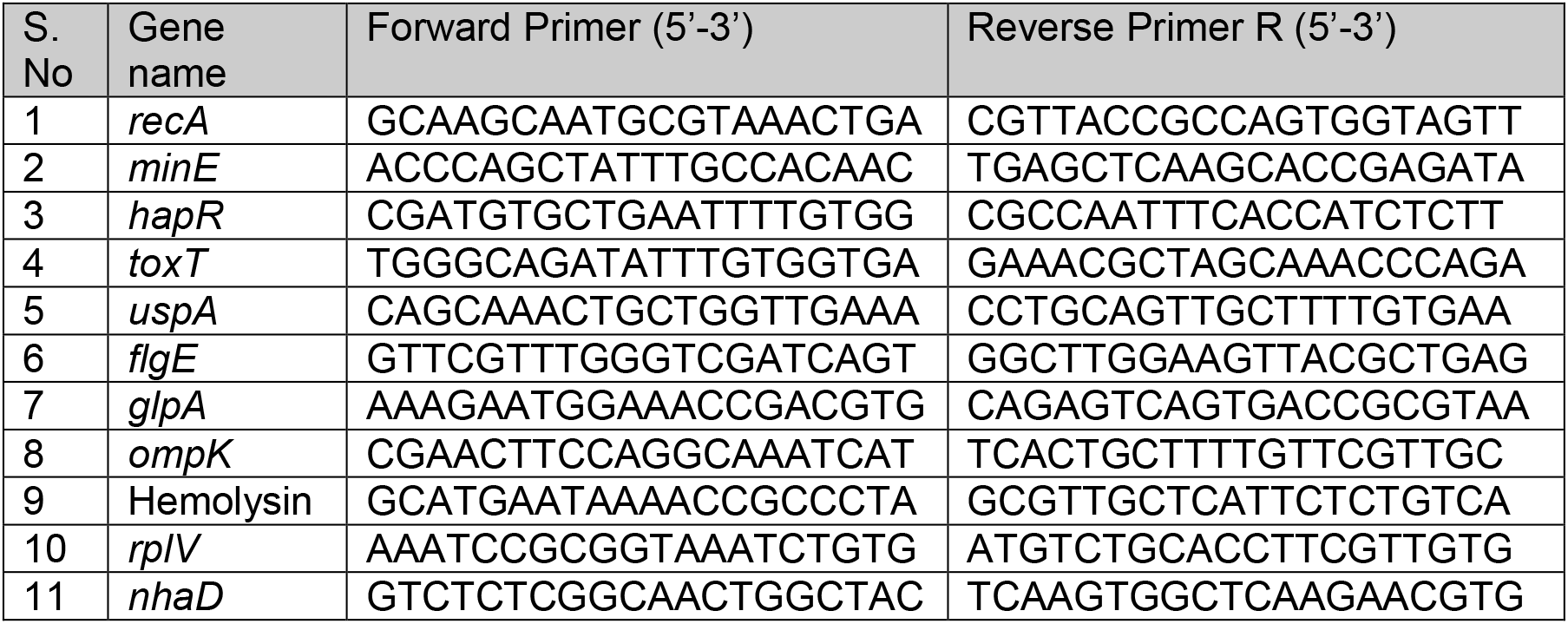
Primers for the genes that were used for gene expression analysis.

## Notes

### Competing Interest Statement

The authors have declared no competing interest.

